# Island-wide population genomic analysis of the coastal shrub, beach naupaka (*Scaevola taccada* L.), on Oʻahu

**DOI:** 10.1101/2025.05.25.655311

**Authors:** Edward V. McAssey, Cassidy Downs, Karolina Heyduk

## Abstract

**Premise:** Plant species face a myriad of challenges including climate change and habitat destruction. On Oʻahu in Hawaiʻi, *Scaevola taccada* L., a coastal dune shrub, is highly impacted by human activities, but is also experiencing environmental threats like sea level rise. Despite its ubiquity across Pacific islands, the extent and distribution of genetic diversity in this species is unclear. Further, it is currently unclear how many times *S. taccada* arrived on Oʻahu.

**Methods:** We surveyed population genomic diversity using ddRAD-seq in eight populations of *S. taccada* on Oʻahu. Using principal components and STRUCTURE analyses, we assessed whether genetic diversity was structured geographically across the island. Additionally, we generated whole genome sequences from five herbarium accessions and used a publicly available genome to construct a haplotype network to infer the number of times *S. taccada* arrived on Oʻahu. We analyzed climate data and leaf thickness measurements to understand how patterns of genetic diversity were correlated with environmental and trait variation.

**Results:** Our analyses pointed to a trend in which two populations in protected areas were both genetically differentiated from, and harbored less genetic diversity than, human dominated areas. Further analysis of genetic structure via a PCA and pairwise F_ST_ indicated a secondary geographic pattern in our dataset by which western beach populations were differentiated from eastern beach populations. The chloroplast haplotype network suggested a single arrival on Oʻahu.

**Conclusions:** Our results demonstrate that *S. taccada* populations in protected natural areas harbored unique genetic variation compared to human dominated beach parks, but that protected populations also had significantly reduced genetic variation compared to other populations.

## Introduction

Islands are an ideal environment to study the dispersal and introduction of plants due to their isolation and relatively small size. Although small, islands contain a greater concentration of endemic species compared to continents (Kier et al., 2009). Isolation, paired with microhabitat variation, can promote speciation, and in some cases, adaptive radiations. However, not all lineages undergo adaptive radiation upon reaching islands (Takayama et al., 2018). In fact, some island species capable of long distance dispersal can have large ranges, such as *Ipomoea pes-caprae* subsp. *Brasiliensis* (L.) van Ooststr., which is found throughout the tropics (Miryeganeh et al., 2014). Another pan tropical species, *Hibiscus tiliaceus* L., has low differentiation among populations throughout its large distribution in part due to seed dispersal via ocean currents (Takayama et al., 2006).

Hawaiʻi is home to approximately 1000 endemic plant species, half of which are at risk of extinction (Sakai et al., 2002). Plant species in Hawaiʻi evolved in numerous habitat types including coastal dunes, lava beds, cliffs, and sub-tropical rainforests. Orographic rain, caused by moisture moving up mountainous terrain, results in sharp environmental gradients in terms of both precipitation and temperature. The habitat variation across Hawai’i has set the stage for a number of adaptive plant radiations including in *Bidens* L. (Helenurm and Ganders, 1985; Knope et al., 2020), *Schiedea* Cham. & Schltdl. (Kapralov and Filatov, 2006), *Plantago* L. (Dunbar-Co et al., 2008), the silversword alliance (Witter and Carr, 1988), and the lobeliads (Givnish et al., 2009).

Other genera in Hawaiʻi, such as *Scaevola* L., inhabit a diverse set of habitats while not undergoing a large adaptive radiation (McKown et al., 2016). *Scaevola* arrived in Hawai’i via at least three independent dispersal events (Howarth et al., 2003). Two of the three dispersals have not undergone cladogenesis, whereas the third dispersal event has resulted in seven endemic species, some of which are homoploid hybrids (Howarth and Baum, 2005). The nine species of *Scaevola* (naupaka in ʻŌlelo Hawaiʻi, Goodeniaceae) span significant environmental gradients in terms of precipitation and elevation that generally correlate with functional traits (McKown et al., 2016). In particular, leaf traits like leaf area, succulence, stomatal density, and vein density were shown to correlate with the environment when comparing all Hawaiian *Scaevola*. Morphological differences between species have long been recognized before colonial documentation of species on the islands. The Hawaiian name for the plant, Naupaka, gets its name from Hawaiian legend, after the princess Naupaka who fell in love with a fisherman (Nishimura, 2016). Their love was forbidden by elders and by the gods, and so Naupaka tore a flower in half and sent the fisherman back to the ocean. Naupaka retreated back into the mountains with her half of the flower. The half-flower shape of *Scaevola* species (Figure 1A) is representative of the two lovers torn apart forever; the legend is also emblematic of the distinct mountain species (naupaka kuahiwi) and the common coastal shrub *Scaevola taccada* Gaertn. (naupaka kahakai).

**Figure 1.**
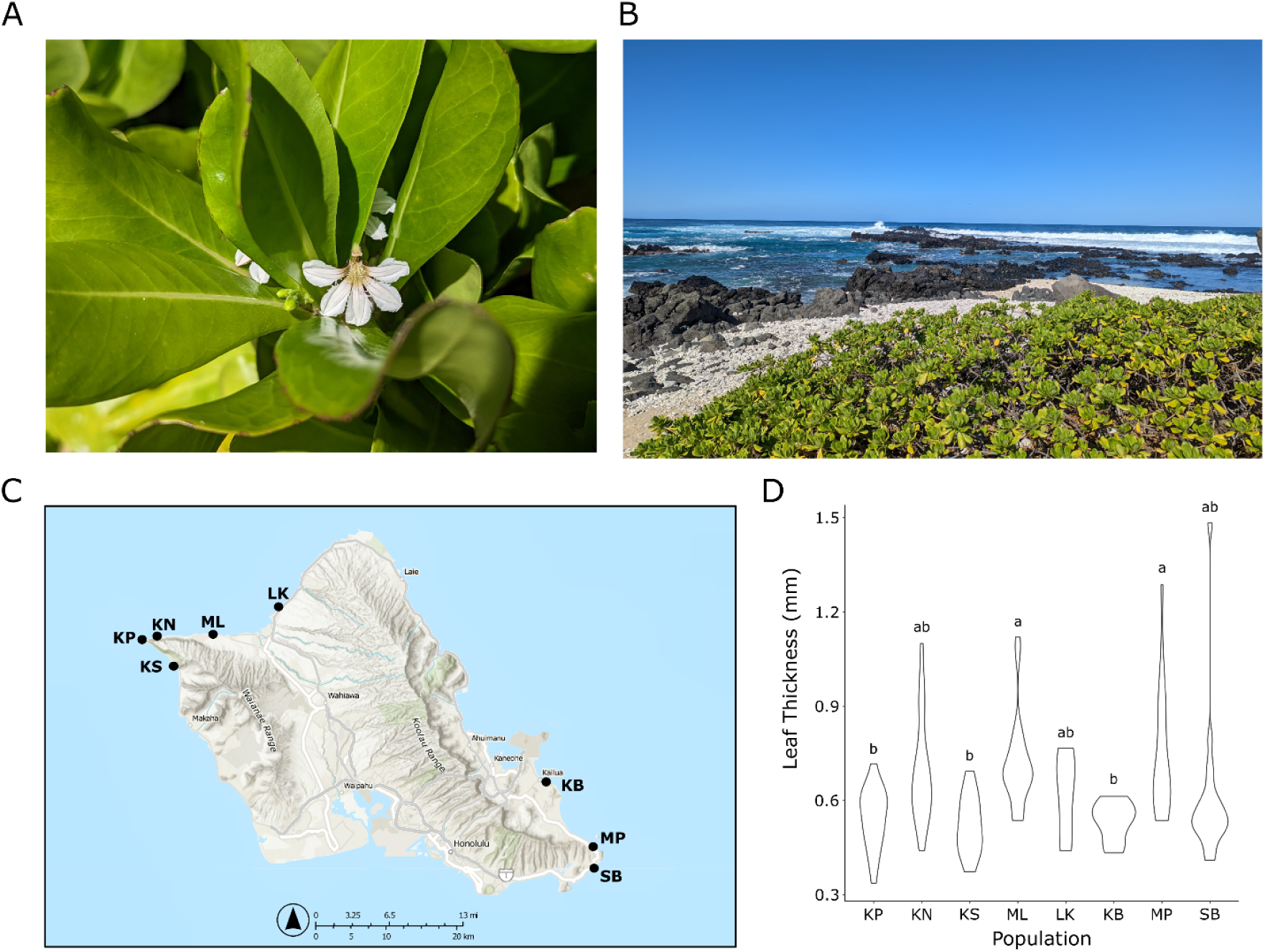
Sampling overview and leaf thickness variation in *S. taccada* A - Flower of *S. taccada*; B - Example dune habitat where *S. taccada* is found (Kaʻena Point); C - Dots correspond to sampling locations on Oʻahu KN - Kaʻena North; KP - Kaʻena Point; KS - Keawaʻula section of Kaʻena Point; KB - Kailua Beach; LK - Laniakea; MK - Makapuʻu; ML - Mokulēʻia; SB - Sandy’s; Map made on Online ArcGIS (Using Esri, NASA, NGA, USGS, TomTom, Garmin, SafeGraph, FAO, METI, EPA, USFWS data); D - Violin plot of leaf thickness variation in each population measured in millimeters

Unlike the *Scaevola* dispersal that produced seven species, *S. taccada* is a widespread coastal dune species in the Pacific basin and has not speciated after reaching Hawaiʻi (Howarth et al., 2003). The fruits of *S. taccada* are capable of floating for at least 180 days (Emura et al., 2014), which promotes its pan-Pacific distribution. However, some populations at higher elevations in south Japan have been shown to have a higher proportion of pulp, which was hypothesized to facilitate dispersal by birds (Emura et al., 2014). While *S. taccada* is a common and abundant species across the greater Pacific basin, it is also an important species of dune ecosystems found in densely human populated islands such as Oʻahu Hawaiʻi (Figure 1B) (Richmond and Mueller-Dombois, 1972), providing habitat for notable species like the Laysan Albatross (*Phoebastria immutabilis*) (Young et al., 2009) and the Hawaiian Yellow-Faced Bee (*Hylaeus anthracinus*) (Graham et al., 2021). Dune ecosystems across the globe are facing threats from climate change, including sea level rise. An analysis of multiple mangrove plant species in southeast asia suggests genetic diversity is being significantly reduced after decades of sea level rise in the region. Despite being adapted to wet environments, multiple mangrove species exhibit low effective population sizes, which is already having an impact on mollusk biodiversity in the area (Guo et al., 2018).

Recently, a genome assembly for an accession of *S. taccada* collected in Dongzhai Harbor, Hainan province, China, was published to enable evolutionary studies in the genus (He et al., 2022; Li et al., 2023). Separately, RAD-seq population genomic analyses were conducted with specimens collected from islands in south Japan to determine levels of gene flow and how gene flow relates to dimorphism in fruit type (cork vs. pulp) (Emura et al., 2022). Genomic analyses indicated that beach *S. taccada* populations exhibited relatively more admixture, presumably due to the relative ease of sea-based dispersal among beach populations, compared to higher elevation populations (Emura et al., 2022). A range-wide study of population genetic variation was conducted in *S. taccada* using nuclear and chloroplast microsatellite markers (Banerjee et al., 2022).The worldwide sampling led to the conclusion that the species originated in Australia before dispersing throughout its contemporary range. This species-wide study precluded more fine scale questions like the introductions to individual islands. For example, only a single accession was studied in Oʻahu (Banerjee et al., 2022), which means the precise introduction history of this populous island in the Pacific remains undescribed.

Our goal in this study was to use genomic data to understand fine scale patterns of population genetic diversity and structure of *S. taccada* within Oʻahu, the third largest island of Hawaiʻi (approximately 1545 km^2^). Oʻahu represents a blend of natural and human dominated environments and *S. taccada* is readily found in both. Relatively few island-wide population genomic studies have occurred on plants on Oʻahu (although there have been previous population genetic studies in comparable study locations by (DeJoode and Wendel, 1992; Cole and Morden, 2021)). Thus, we sought to determine whether the levels and structure of genetic diversity in natural populations on Oʻahu exhibited any significant differences associated with geography and/or human use. We used herbarium accessions from distant Pacific islands to ascertain the number of *S. taccada* introductions to Oʻahu. Finally, we surveyed leaf thickness across populations to determine whether variation in thickness is associated with differences in environmental conditions among populations.

## Methods

### Sampling

*Scaevola taccada* populations were sampled throughout the island of Oʻahu (Figure 1C). Populations were chosen to maximize the likelihood that they represent natural environments, as opposed to areas that are intermixed with housing/resorts that may be planting *S. taccada* as an ornamental plant. In total we sampled eight representative populations: three in east Oʻahu and five in west Oʻahu. These populations varied in terms of human use: six populations within beach parks for recreation and two populations within a state park that is used for fishing and hiking.

Due to its ability to reproduce vegetatively, within each population we sampled plants that were at least approximately 10 meters from one another. Populations varied in terms of the number of plants that we could sample while maintaining a 10 meter buffer between individuals. Population sample sizes ranged from six in Laniakea up to 20 individuals each in Kaʻena North, Mokulēʻia, and Sandy’s Beach with an overall average sampling per population of 15 individuals (Table 1). For each plant, young leaf tissue was sampled into a 15 mL conical tube and stored on ice. Additionally we used digital calipers to measure leaf thickness for the three expanded and mature leaves per plant and averaged the values. Leaf thickness measurements were taken at the widest point of the leaf and care was taken to avoid major veins. The GPS coordinates for all plants were recorded and an herbarium voucher specimen was deposited in the Joseph F. Rock Herbarium at the University of Hawaiʻi at Mānoa for each of the eight populations (Table 1). Tissue samples were stored in a −20 °C freezer until processing.

**Table 1.** Sampling information on Oʻahu.

| Population | N | Latitude | Longitude | Average Annual Rainfall (mm) | Average Driest Month Rainfall (mm) | Herbarium ID |
| --- | --- | --- | --- | --- | --- | --- |
| Ka'ena North (KN) | 20 | 21° 34.773" N | 158° 14.274" W | 768.8 | 22.3 | HAW46034 |
| Ka'ena Point (KP) | 17 | 21° 34.497" N | 158° 16.44" W | 648.9 | 24.8 | HAW46273 |
| Keawa'ula (KS) | 9 | 21° 33.187" N | 158° 14.777" W | 833.6 | 30.4 | HAW46026 |
| Kailua Beach (KB) | 9 | 21° 23.902" N | 157° 43.741" W | 893.2 | 31.5 | HAW46212 |
| Laniakea (LK) | 5 | 21° 37.091" N | 158° 5.145" W | 926.4 | 43.9 | HAW46211 |
| Makapu'u (MP) | 1<br>5 | 21° 18.980"<br>N | 157° 39.835"<br>W | 708.7 | 12.8 | HAW46018 |
| Mokulē'ia (ML) | 1<br>9 | 21° 34.397"<br>N | 158° 11.65" W | 829.8 | 16.7 | HAW46016 |
| Sandy's Beach<br>(SB) | 2<br>0 | 21° 17.148"<br>N | 157° 40.307"<br>W | 698.6 | 12.7 | HAW46021 |

### Phenotypic and environmental analyses

Leaf thickness measurements were averaged for each individual within a population. Population averages were compared via an ANOVA in R (version 4.4.1) and population statistical differences were detected with a Tukey’s HSD test with 95% confidence intervals. Yearly precipitation data and precipitation during the driest months were gathered from an established database (http://rainfall.geography.hawaii.edu/) at the University of Hawaiʻi. Average yearly and driest month precipitation was correlated against average leaf thickness across populations to help determine the extent that environmental variation played a role in establishing trait variation.

### DNA extraction and Library Construction

Frozen leaf tissue was ground under liquid nitrogen with a mortar and pestle, and DNA was isolated with a modified CTAB DNA extraction (Doyle and Doyle, 1987; McAssey et al., 2023). Isolated DNA was quantified using the dsDNA BR Qubit Assay (Invitrogen, Waltham, Massachusetts, USA). DNA was submitted to the Center for Genome Innovation at the University of Connecticut for double digest restriction associated DNA sequencing (ddRAD-seq) library preparation (Peterson et al., 2012).

Samples were digested with *Msp*I and *Pst*I restriction enzymes, and then ligated with barcoded adaptors. The *Msp*I adaptor contained a 10bp universal molecular indicator (UMI) to help identify and resolve PCR duplicates. Ligated products were amplified using Phusion MM (Thermo Fisher Scientific, Waltham, Massachusetts) with the following reaction conditions: 98C for 30 seconds, 11 cycles of 98C for 10 seconds, 65C for 30 seconds, and 72C for 30 seconds, followed by 5 minutes at 72C. PCR product was bead cleaned 1:1, assessed for quality on an Agilent Tapestation D1000 high sensitivity assay, and sequenced on a portion of two separate Illumina NovaSeq 6000 lanes with 100 bp paired reads. De-duplicated and demultiplexed sequencing reads are available on NCBI’s Short Read Archive under BioProject PRJNAXXX.

To understand how many independent arrivals of *S. taccada* occurred on Oʻahu we additionally conducted whole genome sequencing on 12 individuals: five herbarium accessions and seven individuals from our samples collected for ddRAD sequencing (Table S1). The five herbarium samples spanned the equator ranging from 21 S (Cook Island) to 11 N (Taka Island, Republic of the Marshall Islands). DNA extractions for herbarium samples were performed using the same modified CTAB protocol as we did for the extractions from the freshly collected leaves in this study. Whole genome sequencing libraries were made using the KAPA HyperPlus library kit (Roche, Indianapolis, Indiana, USA). Input DNA was quantified via the HS dsDNA Qubit Assay (Invitrogen, Waltham, Massachusetts, USA). DNA samples derived from our fresh leaf tissue DNA extractions were enzymatically fragmented at 37 °C for 15 minutes before undergoing end-repair, A-tailing, and adapter ligation according to the manufacturer’s protocol. Due to the fragmented nature of our herbarium derived DNA, we omitted the enzymatic fragmentation step and began with the end-repair and A-tailing step of the protocol. The barcoded adapters used in this study are Y-stub adapters purchased from the BadDNA lab at the University of Georgia (Glenn et al., 2019). Libraries were 0.8X bead cleaned to remove adapter dimers beforing being amplified for 10 cycles according to the cycling conditions in the KAPA HyperPlus protocol. Amplified libraries were 1X bead cleaned to remove primers and quantified via the HS dsDNA Qubit Assay. Libraries were sequenced on two separate Illumina NovaSeq 6000 lanes with paired end 100 bp reads. Raw sequencing data is available on NCBI’s Short Read Archive under BioProject PRJNAXXX.

### Sequence and population genetic analyses

Stacks v2.64 was used to process and analyze sequence data (Catchen et al., 2013). Within Stacks, the clone_filter script was used to remove PCR clones in our data by looking for exact sequences in paired end reads, which contained the 10 basepair UMI sequence described above. Following clone filtering, Trimmomatic v0.39 was used to remove the UMI sequence, the first ten basepairs of read two (Bolger et al., 2014). The process_radtags script was used to demultiplex reads by barcode and to truncate the ends of read one to 90 basepairs to match the trimmed length of read two. The denovo_map script within Stacks was then used to process all libraries throughout the entire pipeline in an iterative fashion. First, we assembled stacks using -M 2 (max number of mismatches between alleles within an individual) and -m 3 (minimum number read depth to form a stack) and looked to see how many loci were assembled after filtering for loci present in at least 50% of individuals within each of the eight populations. This preliminary analysis results in six libraries missing greater than 25% of the loci established by our filtering criteria. After removing these six libraries, denovo_map was then used to analyze the entire dataset with -M and -n (max number of mismatches between alleles between individuals as they are compared) values of 2 through 5 to see how parameter selection affected stack formation (Catchen et al., 2013).

After determining that the number of loci and SNPs available for population genomic analyses was robust to parameter choice (Table S2), we proceeded with completing all subsequent analyses with the -M and -n values set to 2. The flag -write_single_snp was used in Populations to ensure that no linked variants within a stack were analyzed as they are not independent observations. A minor allele frequency of 0.05 and a maximum observed heterozygosity threshold of 0.7 were both used to retain informative sites. Within Stacks we estimated population genetic parameters like nucleotide diversity, population structure (F_ST_), percent polymorphic loci, observed heterozygosity, and expected heterozygosity. STRUCTURE v2.3.4 was used to further examine the existence of population structure within Oʻahu (Pritchard et al., 2000). Specifically, we used the admixture model with K ranging from 1 to 9. For each value of K we ran 20 replicates with each replicate run consisting of 100,000 burn in and 1,000,000 MCMC iterations. The delta K method (Evanno et al., 2005) was implemented in STRUCTURE Harvester vB.1 (Earl and vonHoldt, 2012) to determine the most likely group of genetic clusters in the dataset. CLUMPP v1.1.2 (Jakobsson and Rosenberg, 2007) and DISTRUCT v1.1 (Rosenberg, 2004) were used to visualize STRUCTURE results. To further verify spatial genetic structure we performed a principal components analysis using the same polymorphisms analyzed in the STRUCTURE analysis above using the R v4.4.1 package SNPRelate (Zheng et al., 2012).

### Chloroplast haplotype network of contemporary and herbarium samples

Raw whole genome sequencing reads were trimmed via Trimmomatic (SLIDINGWINDOW:4:15 LEADING3 TRAILING:3 MINLEN:36; (Bolger et al., 2014)). Due to the small insert size of our fragmented herbarium libraries, we only analyzed read one data and allowed for smaller post-trimmed reads after running Trimmomatic (SLIDINGWINDOW:4:15 LEADING3 TRAILING:3 MINLEN:24). Libraries were mapped to a *S. taccada* chloroplast genome (NCBI record MK397896; (Yao et al., 2019)) using bwa mem v0.7.5a (Li and Durbin, 2009). Two herbarium libraries had poor coverage and were dropped from further analyses. Single nucleotide polymorphisms were called using bcftools v1.9 (Li, 2011). Sites with a depth of at least 3, no heterozygous genotype calls, avoiding the chloroplast inverted repeat regions detected via GeSeq (Tillich et al., 2017), and data present in all 12 individuals were retained for haplotype network construction via Popart (Leigh and Bryant, 2015). A median joining chloroplast haplotype network (epsilon = 0) was constructed from the filtered chloroplast polymorphism data of five herbarium accessions across the Pacific Ocean, seven collections used in the ddRAD sequencing analysis, and the reference chloroplast genome (a cultivated type from Guangdong, China; (Yao et al., 2019)) used for read mapping.

## Results

### Phenotypic and Environmental Variation

Populations had significant variation in leaf thickness (*F* _7,118_ = 4.949, *P* < 0.001), ranging from an average thickness of 0.52 mm at Keawaʻula to 0.75 mm at Makapuʻu (Figure 1D). Populations exhibited very consistent average monthly temperatures for the warmest month of August ranging from 25.41 °C to 25.76 °C. The pattern of annual and summer precipitation was much less constant: average annual precipitation ranged between 648.9 mm at Kaʻena Point to 926.4 mm at Laniakea (Table 1). However, there was no discernible correlation between leaf thickness and annual precipitation. In contrast, when we correlated leaf thickness against the average monthly precipitation for the driest month, we detected a negative, but non-significant, trend (R^2^ = 0.275, *P* > 0.05). Unfortunately, the use of only 8 populations in this study limits the statistical power to dissect this trend further via correlation-based approaches.

### ddRAD-sequencing

An initial run of the Stacks denovo_map pipeline indicated that six samples had > 25% missing loci and were removed from all subsequent analyses. The average number of quality filtered paired reads per individual was 1,389,948 and ranged from 90,567 to 10,305,855. The average number of assembled stacks per individual via uStacks (-M 2; -n 2) was 6,655 and ranged from 2,849 to 19,056. After processing reads through cStacks 54,897 stacks were assembled into the catalog (i.e., were present in at least one individual).

### Population Genetic Diversity

To focus our genetic analysis on shared sites, we required each population to have 50% data present in order to include a locus in the analysis, resulting in 3,230 loci that were analyzed further. The average percent missing loci per individual was 4.65%, and ranged from nearly no missing data (0.46%) to a few individuals with higher rates of missing data (three individuals with >20%; max missing data within an individual was 22.85%). Of the 3,230 filtered loci, 926 contained single nucleotide polymorphisms used for population genetic analyses. The median number of individuals genotyped per population in our dataset was 15.01, with Laniakea having only five, but only six plants were sampled from that population (Table 2). For the 926 analyzed SNPs, within population percent polymorphic loci ranged from 79.26% to 84.02% (Table 2). Populations exhibited significant variation in observed heterozygosity with an average of 0.272 and a range from 0.229 at Kaʻena Point (KP) to 0.325 at Laniakea (LK) (Table 2; *F*_7,7400_ = 20.58, *P* < 0.01). There were also significant differences in the average expected heterozygosity per population with an average of 0.26 and range from 0.219 at Kaʻena Point to 0.277 at Sandy’s Beach (Table 2; *P* < 0.01, *F*_7,7400_ = 11.59). There were some slight differences between populations in the inbreeding coefficient F_IS_ (Table 1; *P* < 0.01; *F*_7,7400_ = 16.96), however, all populations’ F_IS_ values were close to 0 indicating no obvious inbreeding.

**Table 2.**
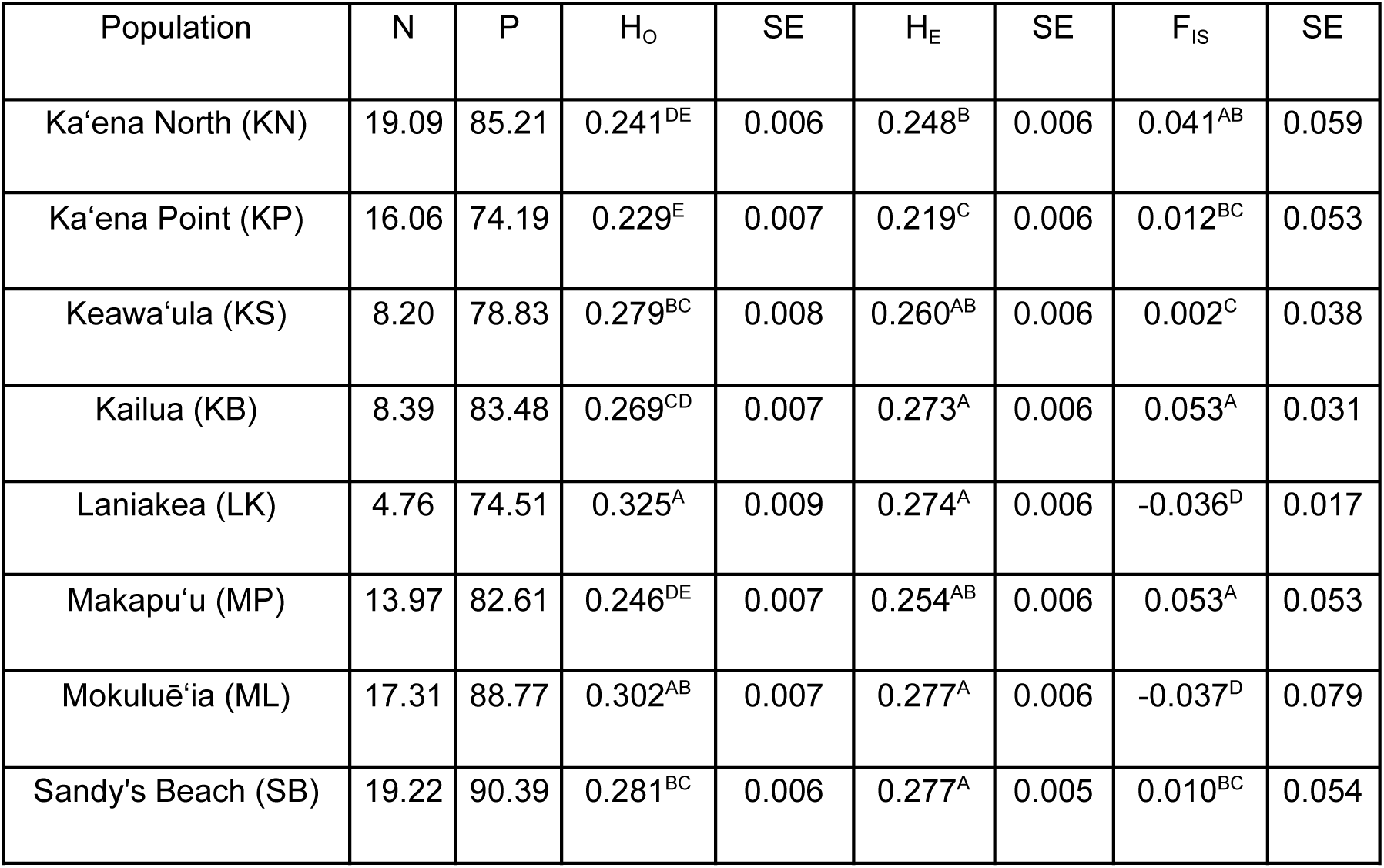
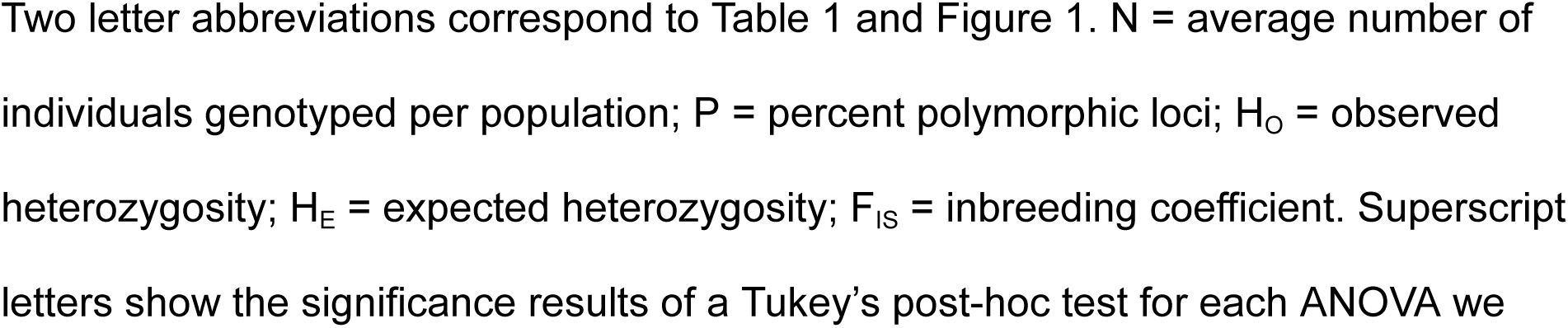
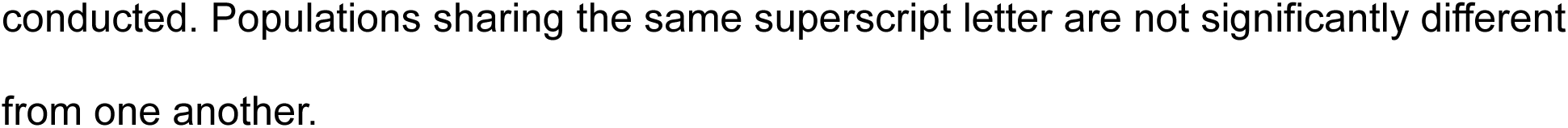
Within population estimates of genetic diversity.

### Population Genetic Structure

Average pairwise F_ST_ values ranged from 0.038 to 0.116 with an average pairwise F_ST_ of 0.0697 (Table 3). The principal component analysis (PCA) of the genotype data identified distinct grouping along PC1, which explains 10.85% of the variance in the dataset. Specifically, the Kaʻena North and Kaʻena Point populations have positive values on PC1 relative to the rest of the dataset (Figure 2). PC2, which explains 5.57% of the variance, is generally separating out the non-Kaʻena North and Kaʻena Point populations by geography. Specifically, negative values on PC2 represent populations on west Oʻahu while positive values are east Oʻahu populations (Figure 2). The PCA results largely correspond to the pairwise F_ST_ results. Both Kaʻena North and Kaʻena Point have consistently greater than average pairwise F_ST_ values when compared to other populations in this dataset. To explore this we removed Kaʻena North and Kaʻena Point from the pairwise F_ST_ analysis and found that the average F_ST_ went down to 0.057, suggesting that those populations are driving the F_ST_ higher (Table 3). This pairwise F_ST_ pattern was highest for Kaʻena Point (average pairwise F_ST_ against all other populations = 0.094), which is consistent with the PCA in which Kaʻena Point individuals are highly differentiated from the six primarily recreational beaches. In our STRUCTURE analysis K=2 was found to be the most likely number of clusters for our Oʻahu sampling (Table S3). One cluster contains Kaʻena Point and Kaʻena North along with a few individuals from the Keawaʻula section of Kaʻena Point (Figure 3). The other cluster contained the other half of Keawaʻula individuals, as well as the remaining populations.

**Figure 2.**
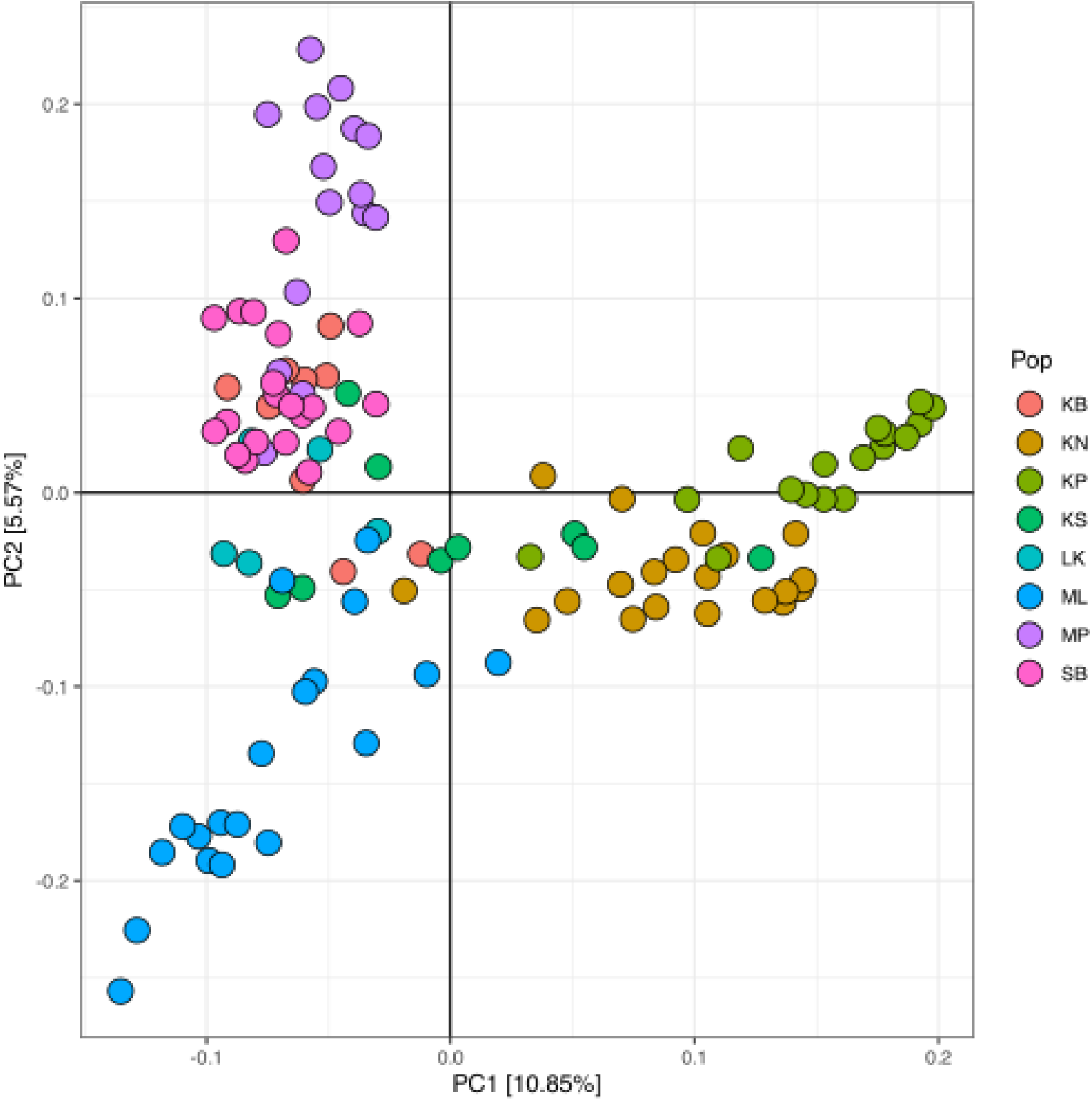
Principal Components Analysis Plot showing the first two principal component axes. PC1 primarily separates protected populations (positive values on PC1) from non-protected populations and explains 10.85% of the variation in the 926 SNP dataset. PC2 explains 5.57% of the variation and generally separates populations geographically with eastern populations having positive values on PC2. Two letter population codes refer to the population labeling in Figure 1 and Table 1.

**Figure 3.**
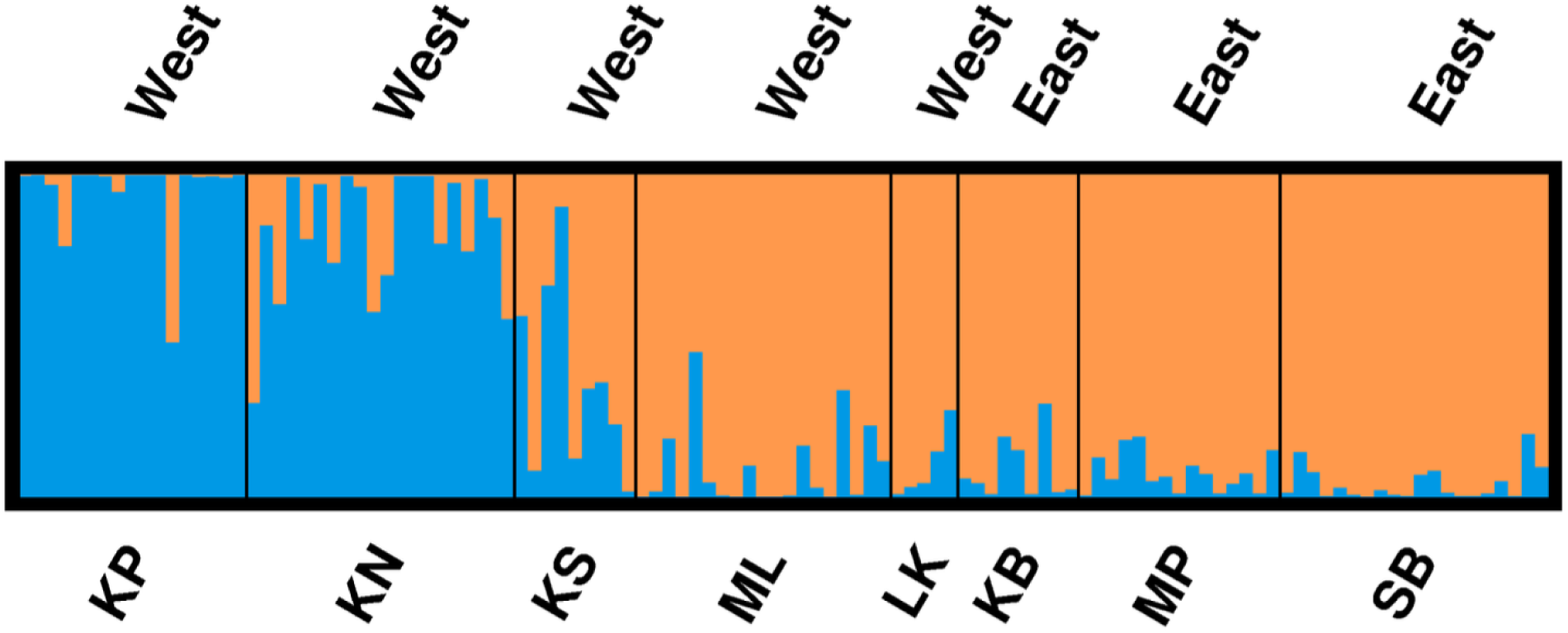
STRUCTURE plot The delta-K method of Evanno et al., 2005 indicated K=2 as being the most likely number of clusters in our dataset. Two letter population codes correspond to Figure 1 and Table 1. East and West labeling refers to the populationʻs position on Oʻahu.

**Table 3.** Pairwise F_ST_ between populations.

|  | Ka'ena Point<br>(KP) | Keawa'ula<br>(KS) | Kailua (KB) | Laniakea<br>(LK) | Makapu'u<br>(MP) | Mokuluē'ia<br>(MK) | Sandy's<br>(SB) |
| --- | --- | --- | --- | --- | --- | --- | --- |
| Ka'ena<br>North (KN) | 0.060 | 0.061 | 0.072 | 0.079 | 0.081 | 0.076 | 0.070 |
| Ka'ena Point<br>(KP) |  | 0.081 | 0.103 | 0.116 | 0.104 | 0.104 | 0.092 |
| Keawa'ula<br>(KS) |  |  | 0.062 | 0.074 | 0.072 | 0.054 | 0.050 |
| Kailua (KB) |  |  |  | 0.056 | 0.060 | 0.055 | 0.038 |
| Laniakea<br>(LK) |  |  |  |  | 0.073 | 0.055 | 0.048 |
| Makapu'u<br>(MP) |  |  |  |  |  | 0.070 | 0.039 |
| Mokuluē'ia<br>(MK) |  |  |  |  |  |  | 0.046 |

To further understand the distribution of population genetic diversity on Oʻahu, we used whole genome skimming (Dodsworth, 2015) to obtain data on samples used in the ddRAD study but also herbarium samples collected throughout the Pacific. 21 chloroplast SNPs were analyzed to form a haplotype network using samples from five herbarium samples from throughout the Pacific Ocean, seven contemporary Oʻahu accessions, and the reference genome (Figure 4). The haplotype network clearly shows most Oʻahu accessions grouping together. The one exception is the accession from Kailua Beach that is more similar to an herbarium sample from the Marshall Islands, than it is to any of the contemporary Oʻahu samples. The reference genome, from China, is closer to all the herbarium accessions than it is to the Oʻahu samples.

**Figure 4.**
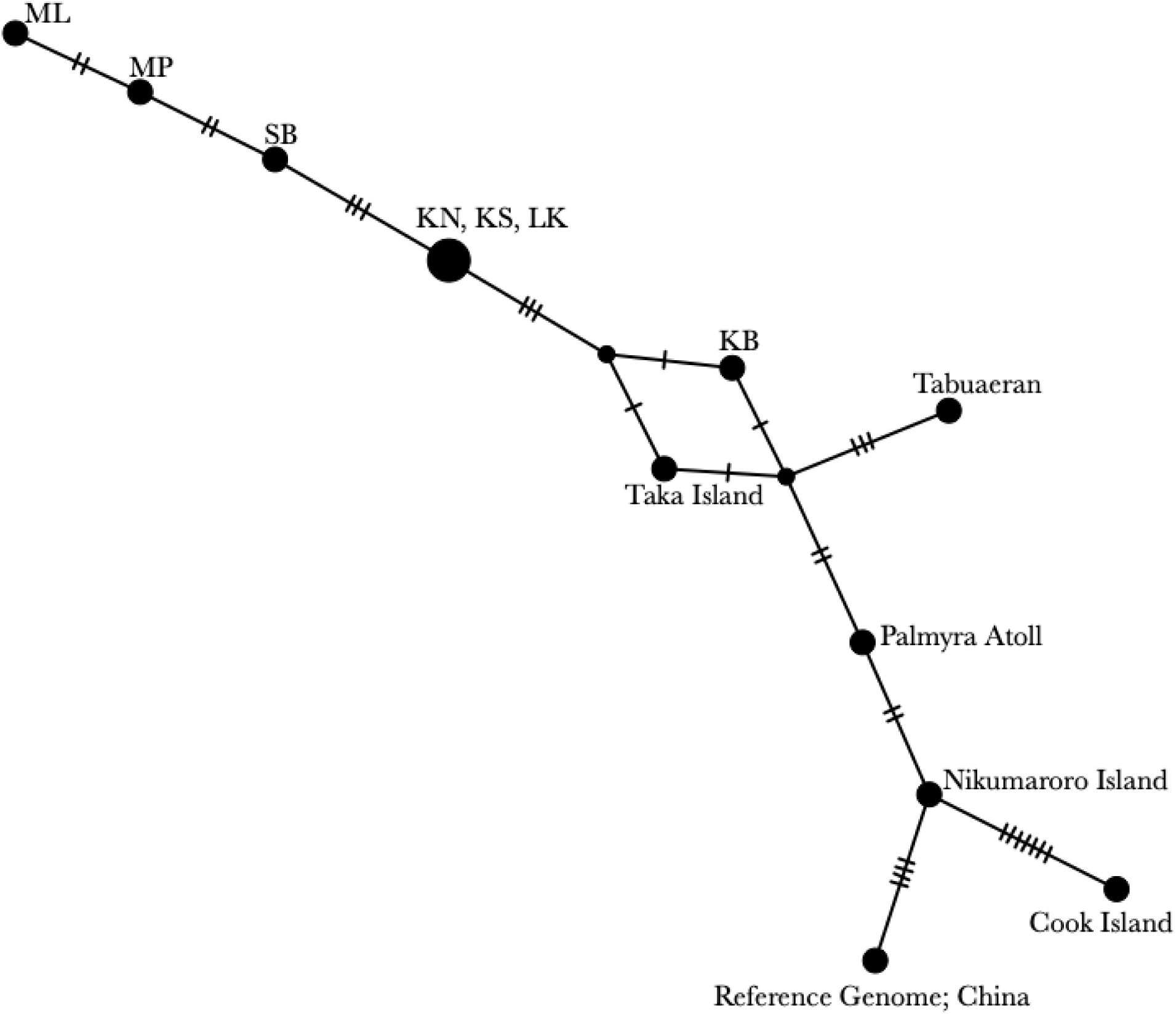
Chloroplast haplotype network Median joining haplotype network. Two letter IDs correspond to Table 1. Full island names represent the five herbarium accessions (Table S1). Hatch marks represent mutations between chloroplast haplotypes.

## Discussion

Our work shows that the ubiquitous dune shrub, *Scaevola taccada*, on the island of Oʻahu harbors significant genetic diversity and shows evidence of population genetic structuring. Paired with the leaf thickness trait analysis, we have shown that a mixture of genetic and phenotypic trait variation exists across a relatively small island. This has important ramifications for the ability of the species to persist under the threat of climate change and sea level rise. The genetic structure of natural populations further indicates protected natural areas harbor distinct, but not more, genetic variation.

Genetic diversity is a prerequisite for adaptation to changing environments. Two populations that are subjected to less human activity, Kaʻena Point and Kaʻena North, both have some of the lower values of genetic diversity, although not statistically different from all populations that experience more human activity. Depending on what part of Oʻahu represents the first area of *S. taccada* occupancy, the relatively lower genetic diversity in Kaʻena could represent a mild founder effect. In general, the presence of genetic diversity within populations suggests that there is in fact a history of sexual reproduction despite their capacity for clonal propagation. Furthermore, the levels of genetic diversity seen throughout Oʻahu suggest that *S. taccada* has not been severely impacted by human mediated or climate change based habitat destruction. Whether this genetic variation is adaptive was not determined in this study. While survival in changing environments can be driven by local adaptation, phenotypic plasticity can also buffer species from environmental change. For example, in a common garden experiment on *Spartina densiflora* Brongn., populations had low genetic diversity, yet still exhibited significant phenotypic variation and plasticity for a number of traits (Castillo et al., 2018). Understanding the extent of local adaptation and the ability for phenotypic plasticity in *S. taccada* would require similar common garden based studies.

We detected significant variation in leaf thickness across populations, but whether these differences are driven by genetic differentiation or environmental conditions is unclear. While the small number of populations precluded a sufficiently statistically powered analysis to detect correlations between leaf thickness and precipitation or temperature, geographically close populations like Kailua Beach and Makapuʻu that were genetically similar (Figure 3) had significant differences in leaf thickness (Figure 1D). The Kailua Beach and Makapuʻu populations also vary in average annual precipitation, with Makapuʻu having 708 mm rain and Kailua Beach having 893 mm rain. Prior work has highlighted the association of functional traits, like leaf succulence, and environmental conditions of *Scaevola* in Hawaiʻi *(McKown et al., 2016).* In *S. taccada*, increased salt exposure led to changes in leaf mass per area (LMA), but not in changes to leaf water content or carbon assimilation rates (Goldstein et al., 1996). A separate study showed increasing δ^13^ values with increased salt exposure (Alpha et al., 1996), suggesting increased water use efficiency. Together the prior work on *S. taccada* suggests investment in leaf mass (or thickness) may be a means by which to limit water loss by decreasing leaf hydraulic conductance; while we did not quantify leaf sap osmolarity of our leaf samples, sea spray and/or inundation variation across populations may be contributing to the significant variation in leaf thickness measured.

Populations on Oʻahu were significantly structured as determined by pairwise F_ST_, STRUCTURE, and PCA. All three methods produced concordant results in which the Kaʻena Point and Kaʻena North populations were significantly different from the other six populations. These populations are more protected (Kaʻena Point is a Natural Area Reserve and Kaʻena North is a State Park) and have larger geographic areas than five of the other populations. The Kaʻena Point area has a long history of human use in the past 200 years (Wahl et al., 2022), including as pasture for railroads, and has served as a military complex. It was only in 1984 that the State Park was established (see the extensive history and documentation of Kaʻena Point found in (Wahl et al., 2022)). The population at Keawaʻula is geographically close to Kaʻena Point and Kaʻena North (being separated by rocky steep terrain) and indeed spans the PC1 axis showing an intermediate position between the protected population cluster (Kaʻena North / Point) and the other five non-protected populations (Figure 2). Mokulēʻia is also relatively close to Kaʻena Point and Kaʻena North, but shows little genetic similarity on PC1 (Figure 2). These results together indicate that geography alone doesn’t explain the patterns of genetic structure. In a species with an ocean dispersing fruit, the ocean currents around the island could explain situations in which the geography doesn’t mirror the population genetic structure. Typically, tradewinds blow out of the northeast towards the southwest when they encounter the Hawaiian islands. These winds impact ocean currents and by extension could dictate seed dispersal. For example, in Hawaiian fish (Counsell et al., 2021) and coral larvae (Storlazzi et al., 2006) currents play a notable role in dispersal dynamics. Current-based dispersal would at least be consistent with the observed differentiation between the east and west Oʻahu populations, although it should be noted that ocean currents are dynamic, change seasonally, and can be disproportionately impacted by large storm systems moving through the area.

There have been a few other studies examining intraspecific genetic diversity of Hawaiian plants across Oʻahu. An endemic Hawaiian cotton, Maʻo or *Gossypium tomentosum* Nutt. ex Seem., was studied throughout the Hawaiian archipelago and across multiple sites on Oʻahu including Makapuʻu and near Kaʻena Point (DeJoode and Wendel, 1992). This study used allozymes to assay genetic variation among few accessions from different locations, but did not sample at the population level and their results are not comparable to our study. However, the authors note that a principal component analysis of their allozyme dataset indicated no geographic relationship with genetic diversity, whereas we saw differentiation within our dataset between those two similar sampling areas. A more recent study on a different Hawaiian endemic species, ʻŌhai or *Sesbania tomentosa* Hook. & Arn., used microsatellite markers to look at genetic diversity across the archipelago including populations in Kaʻena Point, Kaʻena North, as well as an island population just off the coast of Makapuʻu (Cole and Morden, 2021). A STRUCTURE analysis showed that these Oʻahu populations clustered together, indicating an absence of genetic differentiation across Oʻahu. A study on ʻilima or *Sida fallax* Walp. also shared a similar Oʻahu sampling scheme (Pejhanmehr et al., 2024) as our study. Populations sampled from Makapuʻu and Kaʻena Point harbored lower levels of observed heterozygosity (0.1) than what we showed in this study (0.2-0.3; Table 2). The lower genetic diversity in ʻilima was paired with higher genetic differentiation between Makapuʻu and Kaʻena Point (F_ST_ of approximately 0.2), which could reflect differences in pollination and/or seed dispersal (Pejhanmehr et al., 2024). *Scaevola taccada* populations from the southern islands of Japan (Ryukyu, Daito, and Ogasawara) were analyzed to determine levels of gene flow within and among islands (Emura et al., 2022). While the study’s goal involved analyzing a fruit dimorphism, their ddRAD-seq data still provides an interesting comparison point to our study. Genetic differentiation among populations (in their case measured as F_ST_ ^R^) was relatively similar to our findings in Oʻahu (average within island F_ST_ in Japan *S. taccada* was approximately 0.08 compared to 0.07 on Oʻahu [this study]).

The chloroplast haplotype analysis provided no evidence for more than one introduction of *S. taccada* to Oʻahu. However, the sample size for this particular analysis was relatively small, which limited our ability to find multiple introductions. For example, if we had sampled more herbarium accessions from a broader geographic area, we could have encountered accessions that nested within the Oʻahu cluster and thus supported the idea of multiple introductions. However, since the pairwise F_ST_ values between populations in the two different STRUCTURE clusters are relatively low (most are F_ST_ <0.1), the combined nuclear and chloroplast data suggests that a single introduction and subsequent structuring is more likely. Otherwise, the multiple introductions would need to have come from genetically similar source populations. The most obvious immediate source populations that would support that hypothesis are the other Hawaiian islands. A comprehensive phylogeographic study of the whole archipelago could establish whether there are multiple or single introductions from a Hawaiian donor population on another island, and whether intra-island gene flow is ongoing.

In total, this work has highlighted both the diversity and structure of natural populations of *S. taccada* on Oʻahu. While this work showed that there was genetic differentiation, we did not determine the extent to which that variation was adaptive. The fact that protected areas harbored distinct, but less, genetic diversity, shows that population dynamics on a relatively small island can be quite complex. It could suggest that the protected nature of those locations is discouraging terrestrial movement of propagules, thus building up the relatively modest levels of differentiation seen here. Alternatively, these patterns of population structure could be significantly impacted by dispersal patterns from other Hawaiian islands not sampled in this work. This work shows that on a single highly densely settled island in the Pacific Ocean that population genomics can reveal genetic differentiation between high human use and protected areas.

## Acknowledgements

We acknowledge the land this work was conducted on was stolen from the Hawaiian Kingdom, and we thank the Kānaka Maoli for the past and continued stewardship of the Hawaiian islands. We acknowledge various agencies who granted collection permits, including the Hawaiʻi Department of Land and Natural Resources and its Division of State Parks and the Department of Parks and Recreation of the City and County of Honolulu. We also thank the Center for Genome Innovation at UConn and the High Performance Computing Facility at UConn Health. Finally, we thank G. Young Kim for assisting with herbarium specimen vouchering.

## Supplemental Tables

**Table S1.** WGS Herbarium Accession Info.

| Accession number | Sampling location | Nation | Latitude | Longitude |
| --- | --- | --- | --- | --- |
| HAW19140 | Nikumaroro | Kiribati | 4.684439 S | 174.52091 E |
| HAW19145 | Cook Island | Cook Island | 21.204151 S | 159.81871 E |
| HAW19148 | Tabuaeran | Kiribati | 3.871322 N | 159.37441 E |
| HAW19150 | Palmyra Atoll | United States of America | 5.88644 N | 162.08086 E |
| HAW19180 | Taka Island | Republic of the Marshall Islands | 11.112265 N | 169.66733 W |

**Table S2.** Stacks parameter testing.

| M | # loci | # filtered loci | # SNPs |
| --- | --- | --- | --- |
| 2 | 54897 | 3230 | 926 |
| 3 | 54175 | 3217 | 912 |
| 4 | 53785 | 3220 | 942 |
| 5 | 53693 | 3216 | 929 |

M is the Stacks parameter that indicates the maximum number of mismatches allowed between stacks. Assembled loci were filtered so as to only retain loci with at least 50% data present in each of the eight populations.

**Table S3.**
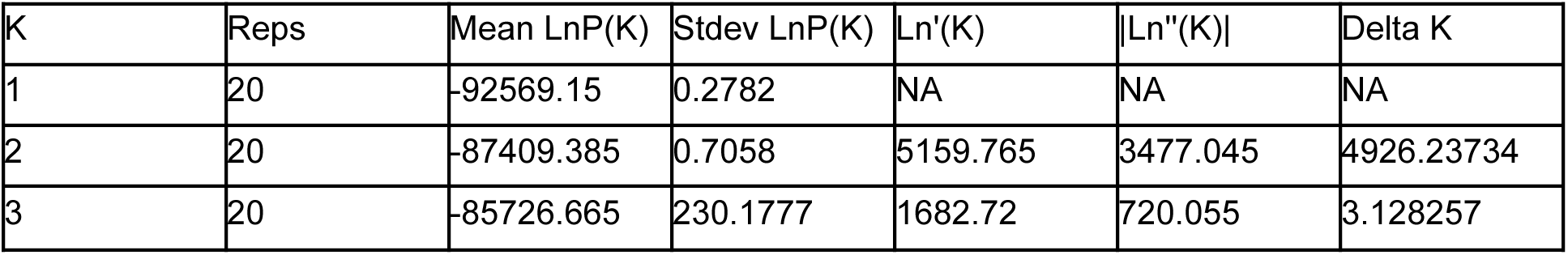
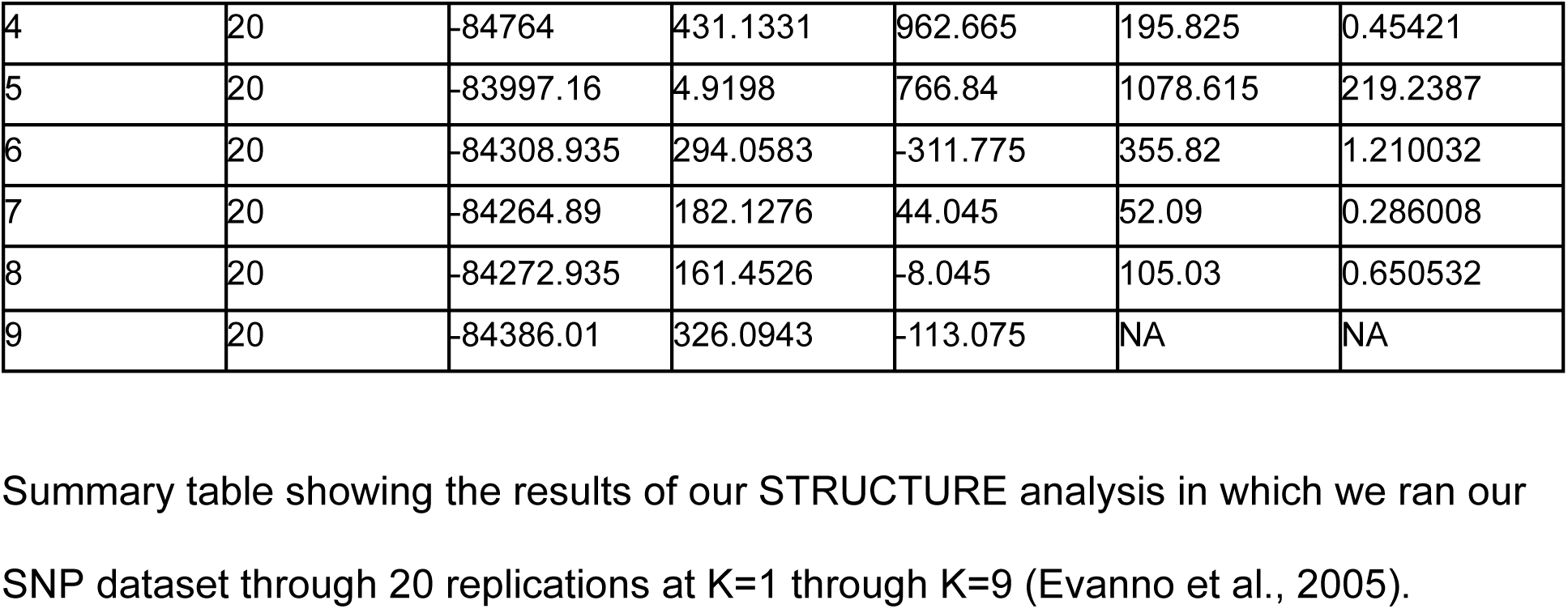
STRUCTURE Delta K analysis.

## References

Alpha, C. G., D. R. Drake, and G. Goldstein. 1996. Morphological and physiological responses of Scaevola sericea (Goodeniaceae) seedlings to salt spray and substrate salinity. American Journal of Botany 83: 86–92.

Banerjee, A. K., H.-D. Wu, W.-X. Guo, W.-L. Ng, W.-X. Li, Y. Ma, H. Feng, and Y.-L. Huang. 2022. Deciphering the global phylogeography of a coastal shrub ( *Scaevola taccada* ) reveals the influence of multiple forces on contemporary population structure. Journal of Systematics and Evolution 60: 809–823.

Bolger, A. M., M. Lohse, and B. Usadel. 2014. Trimmomatic: a flexible trimmer for Illumina sequence data. Bioinformatics 30: 2114–2120.

Castillo, J. M., B. Gallego-Tévar, E. Figueroa, B. J. Grewell, D. Vallet, H. Rousseau, J. Keller, et al. 2018. Low genetic diversity contrasts with high phenotypic variability in heptaploid Spartina densiflora populations invading the Pacific coast of North America. Ecology and Evolution 8: 4992–5007.

Catchen, J., P. A. Hohenlohe, S. Bassham, A. Amores, and W. A. Cresko. 2013. Stacks: an analysis tool set for population genomics. Molecular Ecology 22: 3124–3140.

Cole, D. M., and C. W. Morden. 2021. Population Divergence and Evolution of the Hawaiian Endemic Sesbania tomentosa (Fabaceae). Pacific Science 75.

Counsell, C. W. W., R. R. Coleman, S. S. Lal, B. W. Bowen, E. C. Franklin, A. B. Neuheimer, B. S. Powell, et al. 2021. Opening the black box: interdisciplinary analysis of larval dispersal for a coral reef fish. Marine Ecology Progress Series 684: 117–132.

DeJoode, D. R., and J. F. Wendel. 1992. Genetic diversity and origin of the Hawaiian islands cotton, Gossypium tomentosum. American Journal of Botany 79: 1311–1319.

Dodsworth, S. 2015. Genome skimming for next-generation biodiversity analysis. Trends in Plant Science 20: 525–527.

Doyle, J. J., and J. L. Doyle. 1987. A rapid DNA isolation procedure for small quantities of fresh leaf tissue. Phytochemical Bulletin.

Dunbar-Co, S., A. M. Wieczorek, and C. W. Morden. 2008. Molecular phylogeny and adaptive radiation of the endemic Hawaiian Plantago species (Plantaginaceae). American Journal of Botany 95: 1177–1188.

Earl, D. A., and B. M. vonHoldt. 2012. STRUCTURE HARVESTER: a website and program for visualizing STRUCTURE output and implementing the Evanno method. Conservation Genetics Resources 4: 359–361.

Emura, N., T. Denda, M. Sakai, and K. Ueda. 2014. Dimorphism of the seed-dispersing organ in a pantropical coastal plant, Scaevola taccada: heterogeneous population structures across islands. Ecological Research 29: 733–740.

Emura, N., T. Muranaka, T. Iwasaki, M. N. Honjo, A. J. Nagano, Y. Isagi, and H. Kudoh. 2022. Effects of fruit dimorphism on genetic structure and gene flow in the coastal shrub Scaevola taccada. Annals of Botany 130: 1029–1040.

Evanno, G., S. Regnaut, and J. Goudet. 2005. Detecting the number of clusters of individuals using the software STRUCTURE: a simulation study. Molecular Ecology 14: 2611–2620.

Givnish, T. J., K. C. Millam, A. R. Mast, T. B. Paterson, T. J. Theim, A. L. Hipp, J. M. Henss, et al. 2009. Origin, adaptive radiation and diversification of the Hawaiian lobeliads (Asterales: Campanulaceae). Proceedings of the Royal Society B 276: 407–416.

Glenn, T., R. Nilsen, T. J. Kieran, J. Sanders, N. Bayona-Vásquez, J. W. Finger, T. W. Pierson, et al. 2019. Adapterama I: universal stubs and primers for 384 unique dual-indexed or 147,456 combinatorially-indexed Illumina libraries (iTru & iNext). PeerJ 7.

Goldstein, G., D. R. Drake, C. Alpha, P. Melcher, J. Heraux, and A. Azocar. 1996. Growth and photosynthetic responses of Scaevola sericea, A Hawaiian coastal shrub, to substrate salinity and salt spray. International Journal of Plant Sciences 157: 171–179.

Graham, J. R., J. W. Campbell, S. Plentovich, and C. B. A. King. 2021. Nest architecture of an endangered Hawaiian yellow-faced bee, Hylaeus anthracinus (Hymenoptera: Colletidae) and potential nest-site competition from three introduced solitary Bees1. Pacific Science 75: 361–370.

Guo, Z., X. Li, Z. He, Y. Yang, W. Wang, C. Zhong, A. J. Greenberg, et al. 2018. Extremely low genetic diversity across mangrove taxa reflects past sea level changes and hints at poor future responses. Global Change Biology 24: 1741–1748.

Helenurm, K., and F. R. Ganders. 1985. Adaptive radiation and genetic differentiation in Hawaiian Bidens. Evolution 39: 753–765.

He, Z., X. Feng, Q. Chen, L. Li, S. Li, K. Han, Z. Guo, et al. 2022. Evolution of coastal forests based on a full set of mangrove genomes. Nature Ecology & Evolution 6: 738–749.

Howarth, D. G., and D. A. Baum. 2005. Genealogical evidence of homoploid hybrid speciation in an adaptive radiation of Scaevola (Goodeniaceae) in the Hawaiian Islands. Evolution 59: 948–961.

Howarth, D. G., M. H. G. Gustafsson, D. A. Baum, and T. J. Motley. 2003. Phylogenetics of the genus Scaevola (Goodeniaceae): implication for dispersal patterns across the Pacific Basin and colonization of the Hawaiian Islands. American Journal of Botany 90: 915–923.

Jakobsson, M., and N. Rosenberg. 2007. CLUMPP: a cluster matching and permutation program for dealing with label switching and multimodality in analysis of population structure. Bioinformatics 23: 1801–1806.

Kapralov, M. V., and D. A. Filatov. 2006. Molecular adaptation during adaptive radiation in the Hawaiian endemic genus Schiedea. PloS One 1: e8.

Kier, G., H. Kreft, T. M. Lee, W. Jetz, P. L. Ibisch, C. Nowicki, J. Mutke, and W. Barthlott. 2009. A global assessment of endemism and species richness across island and mainland regions. Proceedings of the National Academy of Sciences, USA 106: 9322–9327.

Knope, M. L., M. R. Bellinger, E. M. Datlof, T. J. Gallaher, and M. A. Johnson. 2020. Insights into the Evolutionary History of the Hawaiian Bidens (Asteraceae) Adaptive Radiation Revealed Through Phylogenomics. Journal of Heredity 111: 119–137.

Leigh, J. W., and D. Bryant. 2015. Popart: Full-feature software for haplotype network construction. Methods in Ecology and Evolution 6: 1110–1116.

Li, H. 2011. A statistical framework for SNP calling, mutation discovery, association mapping and population genetical parameter estimation from sequencing data. Bioinformatics 27: 2987–2993.

Li, H., and R. Durbin. 2009. Fast and accurate short read alignment with Burrows-Wheeler transform. Bioinformatics 25: 1754–1760.

Li, S., X. Mao, Z. He, S. Xu, Z. Guo, and S. Shi. 2023. Chromosomal-scale genome assemblies of two coastal plant species, Scaevola taccada and S. hainanensis-insight into adaptation outside of the common range. International Journal of Molecular Sciences 24: 7355.

McAssey, E. V., C. Downs, M. Yorkston, C. W. Morden, and K. Heyduk. 2023. A comparison of freezer-stored DNA and herbarium tissue samples for chloroplast assembly and genome skimming. Applications in Plant Sciences.

McKown, A. D., M. E. Akamine, and L. Sack. 2016. Trait convergence and diversification arising from a complex evolutionary history in Hawaiian species of Scaevola. Oecologia 181: 1083–1100.

Miryeganeh, M., K. Takayama, Y. Tateishi, and T. Kajita. 2014. Long-distance dispersal by sea-drifted seeds has maintained the global distribution of Ipomoea pes-caprae subsp. brasiliensis (Convolvulaceae). PloS One 9: e91836.

Nishimura, M. 2016. Legend of naupaka comes to life in Kaka’ako murals. Kamehameha Schools. Website https://www.ksbe.edu/article/legend-of-naupaka-comes-to-life-in-kakaako-murals [accessed 5 February 2025].

Pejhanmehr, M., M. B. Kantar, M. Yorkston, and C. W. Morden. 2024. Population genetics of Sida fallax Walp. (Malvaceae) in the Hawaiian Islands. Frontiers in Plant Science 15: 1304078.

Peterson, B. K., J. N. Weber, E. H. Kay, H. S. Fisher, and H. E. Hoekstra. 2012. Double digest RADseq: an inexpensive method for de novo SNP discovery and genotyping in model and non-model species. PloS One 7: e37135.

Pritchard, J. K., M. Stephens, and P. Donnelly. 2000. Inference of population structure using multilocus genotype data. Genetics 155: 945–959.

Richmond, T. de A., and D. Mueller-Dombois. 1972. Coastline ecosystems on Oahu, Hawaii. Plant Ecology 25: 367–400.

Rosenberg, N. A. 2004. Distruct: A program for the graphical display of population structure. Molecular Ecology Notes 4: 137–138.

Sakai, A. K., W. L. Wagner, and L. A. Mehrhoff. 2002. Patterns of endangerment in the hawaiian flora. Systematic Biology 51: 276–302.

Storlazzi, C. D., E. K. Brown, and M. E. Field. 2006. The application of acoustic Doppler current profilers to measure the timing and patterns of coral larval dispersal. Coral Reefs 25: 369–381.

Takayama, K., D. J. Crawford, P. López-Sepúlveda, J. Greimler, and T. F. Stuessy. 2018. Factors driving adaptive radiation in plants of oceanic islands: a case study from the Juan Fernández Archipelago. Journal of Plant Research 131: 469–485.

Takayama, K., T. Kajita, J. Murata, and Y. Tateishi. 2006. Phylogeography and genetic structure of *Hibiscus tiliaceus*— speciation of a pantropical plant with sea-drifted seeds. Molecular Ecology 15: 2871–2881.

Tillich, M., P. Lehwark, T. Pellizzer, E. S. Ulbricht-Jones, A. Fischer, R. Bock, and S. Greiner. 2017. GeSeq--versatile and accurate annotation of organelle genomes. Nucleic Acids Research 45: W6–W11.

Wahl, M., T. Luthy, and K. ʻāhiki Solis. 2022. Archival research documenting the cultural and natural resources of Kaʻena point in support of a national heritage area designation.

Witter, M. S., and G. D. Carr. 1988. Adaptive radiation and genetic differentiation in the Hawaiian silversword Alliance (Compositae: Madiinae). Evolution 42: 1278–1287.

Yao, G., J.-J. Jin, H.-T. Li, J.-B. Yang, V. S. Mandala, M. Croley, R. Mostow, et al. 2019. Plastid phylogenomic insights into the evolution of Caryophyllales. Molecular Phylogenetics and Evolution 134: 74–86.

Young, L. C., E. A. Vanderwerf, D. G. Smith, J. Polhemus, N. Swenson, C. Swenson, B. R. Liesemeyer, et al. 2009. Demography and natural history of laysan albatross on Oahu, Hawaii. The Wilson Journal of Ornithology 121: 722–729.

Zheng, X., D. Levine, J. Shen, S. M. Gogarten, C. Laurie, and B. S. Weir. 2012. A high-performance computing toolset for relatedness and principal component analysis of SNP data. Bioinformatics 28: 3326–3328.

